# Homeostatic regulation of intrinsic lipid curvature in eukaryotic cells

**DOI:** 10.1101/2025.10.13.682232

**Authors:** Daniel Milshteyn, Jacob R. Winnikoff, Elida Kocharian, Aaron M. Armando, Edward A. Dennis, Peter R. Girguis, Itay Budin

## Abstract

Cell membranes are composed of both bilayer-supporting and non-bilayer phospholipids, with the latter’s negative intrinsic curvature aiding in membrane trafficking and the dynamics of membrane proteins. Phospholipid metabolism has long been recognized to maintain membrane fluidity, but whether it also acts to maintain the function of high-curvature lipids is not resolved. Here, we find that cells grown under hydrostatic pressure – used to artificially reduce lipid curvature – maintain lipidome curvature through metabolic acclimation. We first observed that manipulation of the lipidome curvature via the phosphatidylethanolamine (PE) to phosphatidylcholine (PC) ratio affects high-pressure growth and viability of yeast independently of membrane fluidity. In wild-type cells, X-ray scattering measurements revealed an increased propensity for lipid extracts to form non-lamellar phases after extended pressure incubations. Unexpectedly, this change in phase behavior was not due to increased levels of PE, but of phosphatidylinositol (PI), the only major phospholipid class whose curvature had not been previously characterized. We found that PI is a non-bilayer lipid, with a negative curvature intermediate to that of PE and PC. Accounting for PI, mean lipidome curvature was defended in response to pressure by two distantly related yeasts. Lipidome curvature also responded to pressure in a human cancer cell line through ether phospholipid metabolism and chain remodeling, but not in bacterial cells. These findings indicate that eukaryotic phospholipid metabolism uses diverse mechanisms to maintain curvature frustration in cell membranes.

## Introduction

The chemistry of cellular lipids determines their shape and propensity to pack into both bilayer and non-bilayer structures. Phospholipids that are cylindrical in shape, like phosphatidylcholine (PC), pack readily into lamellar phases (bilayers). Since they form leaflets that are mechanically unstressed with little tendency to curve, such lipids are considered to have intrinsic lipid curvature (c_0_) near zero. Other abundant lipids, like phosphatidylethanolamine (PE), have a smaller cross-section at the headgroup than in the hydrocarbon region, giving them a conical shape. In isolation, such phospholipids form non-bilayer structures, like the inverse hexagonal (H_II_) or bicontinuous cubic (Q_II_) phases, in which lipid monolayers curve towards the head groups (1) (Fig. 1A). This is defined by convention as negative curvature, so such lipids have c_0_ < 0. They are referred to herein as high-curvature lipids, since we primarily consider species with c_0_ ≤ 0. When incorporated into flat membranes, high-curvature lipids impose packing frustration also known as curvature elastic stress (2). In some compartments, like the inner mitochondrial membrane, non-bilayer phospholipids make up the majority of the membrane (3). The observation that cells synthesize diverse phospholipids that destabilize bilayer structure seems antithetical to the classical model of cell membranes as robust barrier structures. High-curvature lipids appear to balance this barrier function with a requirement for membrane dynamics. They modulate membrane protein conformations through changes the bilayer’s internal stresses (4, 5), generate packing defects for interactions with soluble proteins (6), and facilitate otherwise unfavorable structural intermediates in membrane fission and fusion processes (7, 8). The relative amount of lipids that stabilize versus frustrate membranes has been proposed as a metabolic lever for keeping these structures robust yet dynamic (9).

**Fig. 1:**
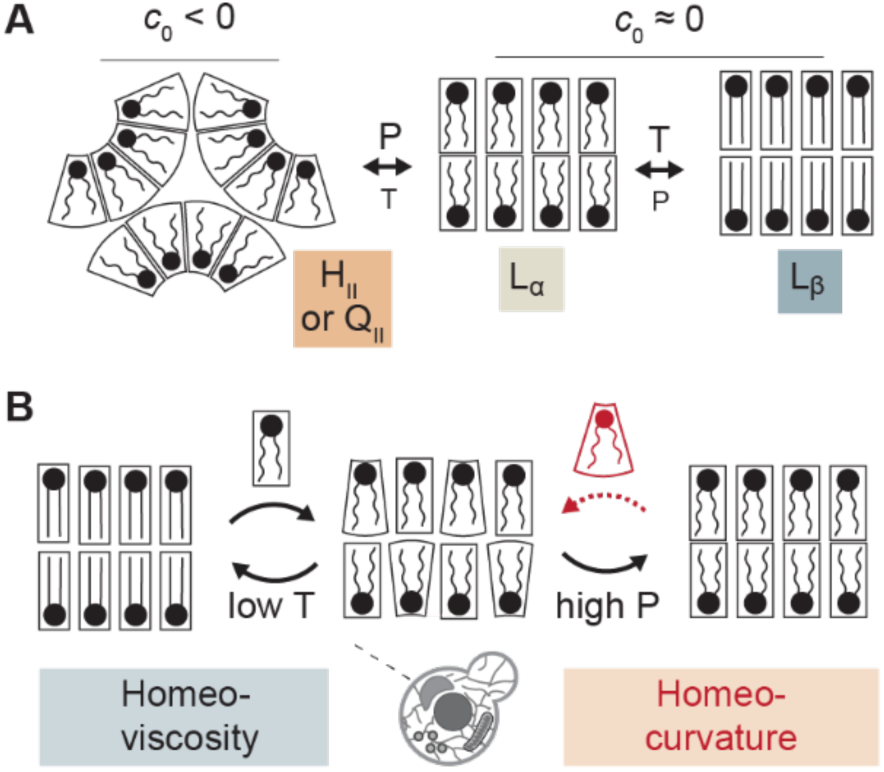
Homeostasis of cell membranes based on membrane fluidity and lipid curvature. **(*A*)** Cell membrane lipid building blocks can have curvatures (*c0*) close to zero, in which case they form lamellar phases like Lα (fluid) and Lβ (solid), but often feature negative *c0* that promote non-lamellar phases like HII or QII. Temperature (T) and pressure (P) can both shift lipids between these phases, but T is a stronger modulator of fluidity (Lα to Lβ) and P of curvature (HII/QII to Lα). **(*B*)** Homeoviscosity – maintenance of fluidity (left) – is well-established as a cellular response to temperature. Mechanisms for homeocurvature – maintenance of lipid shape (right) – are less explored but would be triggered in response to pressure.

Cell metabolism has long been known to maintain key properties of membranes by altering lipid stoichiometry in response to environmental changes. Classically, this homeostasis has been understood through the capacity of a wide range of organisms to maintain the fluidity of their membranes in response to temperature. Membranes contain mixtures of low- and high-melting temperature (T_m_) lipids, which control membrane fluidity and are required for mechanisms of lateral organization. Low temperatures or increases in high-T_m_ lipids can drive fluid lamellar-phase (L_α_) bilayers to a solid or gel-like phase (L_β_) (10, 11). Cells control the fluidity of their bilayers and avoid L_β_ by modulating unsaturation of phospholipid radyl chains (12, 13), the hydrogen bonding capacity of their head groups through changes in the PE/PC ratio (14), or the relative amount of other lipid classes, like sterols (15). In some systems, sensors for fluidity-related properties have been identified, which respond to changes in bilayer packing or thickness and transcriptionally regulate unsaturated fatty acid metabolism (16, 17). While these responses – generally termed homeoviscous adaptation or acclimation – are activated during changes in growth temperature, they are also required for general optimization of fluidity-mediated processes like respiration (18).

Several lines of evidence have suggested that cells utilize homeostatic mechanisms for maintaining curvature stress analogous to those that maintain fluidity (2, 9, 19–21). However, a consistent challenge in their interpretation is that experimental levers observed to cause changes in non-bilayer lipids – which include temperature or mutations in phospholipid metabolism – have pleiotropic effects on lipid properties. For example, while low temperatures reduce curvature by restricting the volume of phospholipid chains (19), they more strongly reduce membrane fluidity and promote gel phases, as described below. Similarly, restriction of PC synthesis promotes negative curvature and drives changes in PE metabolism (22), but loss of PC also inhibits membrane fluidity due to its low T_m_ (14). Thus, it is unclear whether previously observed lipid changes were truly serving to maintain lipid curvature or were instead secondary effects of responses to membrane fluidity and associated properties.

Recent work highlights hydrostatic pressure as an environmental variable that acts disproportionately on lipid intrinsic curvature. Among ctenophores (comb jellies), animals specialized for high pressure exhibit phospholipid chemistry that defends negative lipid curvature and access to non-bilayer phases (homeocurvature adaptation) (23). Pressure drives biomolecules into lower-volume conformations and has an especially large effect on lipids due to the high compressibility of their hydrocarbon chains. Pressure has a stronger effect on membrane curvature than on fluidity because phospholipid compressibility is anisotropic: the hydrocarbon chain volume is more pressure-sensitive than that of head groups. In model membranes, the temperature-pressure equivalency (dT/dP) of the curvature-mediated L_α_ → H_II_ transition is 0.05 K/bar (24, 25), e.g. a pressure of 400 bar corresponds to a temperature drop of ∼20 °C. In contrast, the dT/dP for the L_α_ → L_β_ transition is 3-fold smaller, so fluidity is dominated by temperature and weakly affected by pressure. In ctenophores, adaptation to high-pressure involves synthesis of the high-curvature plasmenyl PE (P-PE), as well as incorporation of longer chains across all phospholipids (23, 26). In vitro, pressure inhibits processes like membrane fusion (23) that are dependent on lipid intrinsic curvature (27).

We hypothesized that if cells sense and regulate intrinsic curvature directly, then they would alter their lipid chemistry in response to high-pressure treatment that directly reduces it. Since terrestrial organisms are unlikely to have evolved *bona fide* hydrostatic pressure sensors, such acclimation would be driven by changes in membrane packing. In this way, high pressure could be utilized to delineate mechanisms of homeocurvature acclimation from other modes of membrane homeostasis in non-pressure adapted cells (Fig. 1B). Here we use this approach to identify an endogenous capacity of phospholipid metabolism to modulate lipidome curvature in both yeast and a human cancer cell line.

## Results

### Lower PE content reduces stability of non-lamellar phases and hydrostatic pressure tolerance

High pressures decrease the curvature of non-bilayer lipids (24), challenging cells’ ability to access non-lamellar membrane topologies. As PC and PE are the most abundant lamellar and non-lamellar lipids, respectively, in eukaryotic lipidomes, we chose to screen the effects of varying PE/PC ratios on fitness of yeast grown at pressure. The *PSD1* and *PEM1/2* genes in *Saccharomyces cerevisiae* (W303) are predominantly responsible for PE and PC biosynthesis in the CDP-DAG pathway (28). Alternative pathways for PE and PC synthesis channel through the CDP-choline or -ethanolamine branches of the Kennedy pathway and require choline or ethanolamine supplementation, offering an opportunity for rescue controls in defined growth media. Phospholipid profiles of wild-type (WT), *pem2*Δ, *psd1*Δ, and supplementation of 2 mM choline to *psd1*Δ (*psd1Δ* + CHO) were determined by liquid chromatography-tandem mass spectrometry (LC-MS^2^) (Fig 2A). Disruption of PE methylation in *pem2*Δ resulted in an almost complete loss of PC. In the *psd1*Δ mutant, PE was depleted by >75% and partially rescued when supplemented with choline. Along with reduced PE or PC in the respective knockout strains, an increase in PS was observed in strains with lower PE levels, as PS is the substrate for PE-generating Psd1. A notable increase in PI was also observed in *pem2*Δ and *psd1*Δ cells.

**Fig. 2:**
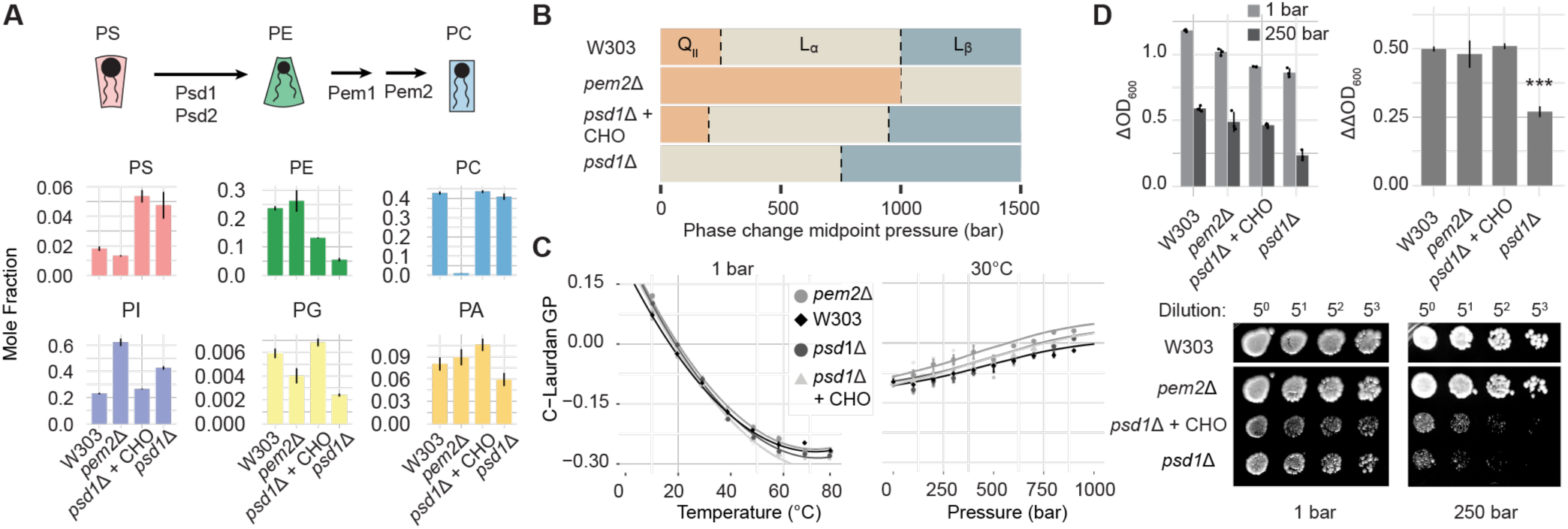
Reduced access to non-lamellar phases decreases high-pressure fitness. *(**A**)* Phospholipid levels of wild-type (W303) yeast and mutants with knockouts in PC biosynthesis (*pem2*Δ), PE biosynthesis (*psd1Δ*), and *psd1Δ* with supplementation of 2 mM choline as a partial rescue (*psd1*Δ + CHO). Error bars indicate SEM from replicates of individually grown cultures *N* = 3. *(**B**)* Pressure-dependent phases of extracted polar lipids for each strain as determined by HPSAXS. Cells were grown at 1 bar and 30°C; measurements were performed at 85°C. Scattering profiles shown in Supp. Fig. 1. Cells lacking Psd1 show reduced prevalence of the non-bilayer cubic phase (QII), while those lacking Pem2 show a stabler QII. *(**C**)* C-Laurdan GP values from polar lipid extracts of strains in HPFS experiments across a temperature range (5-80 °C) at 1 bar and a pressure range (1-1000 bar) at 30°C, respectively. Increasing GP values indicate a lower membrane fluidity, but the strains showed similar behaviors. Full P/T profiles are shown in Supp. Fig. 2. *(**D**)* Growth (top) and survival (bottom) of PE/PC variant strains after 1 or 250 bar pressure incubation. Cells lacking Psd1 show reduced growth (*psd1*Δ) and survival (*psd1*Δ, *psd1*Δ + CHO). Error bars indicate SEM from *N* = 3 replicates of individually grown cultures. *** *P* < 0.001, 1-way ANOVA with Dunnet’s test of *psd1*Δ compared against W303.

We next asked how changes in PE and PC affect the biophysical properties of yeast lipids. To characterize access to non-lamellar lipid phases, we measured high-pressure small angle X-ray scattering (HPSAXS) of rehydrated polar lipid extracts from cultures grown at 1 bar. Samples were equilibrated to 85°C to ensure the formation of non-lamellar phases before isothermally increasing pressure (Fig 2B). At 1 bar, scattering profiles for all strains except *psd1*Δ displayed a sharp primary peak near 0.05 Å^-1^ whose intensity decreased with pressure; we interpreted this as a non-lamellar Q_II_ phase with repeat spacing of 2π/0.05 Å^-1^ ≈ 125 Å (Supp. Fig. 1). As pressure increased, peaks with 1:2:3:4… q-spacing emerged, representing the diffraction signature of fluid (L_α_) or solid (L_β_) lamellar membranes. The sensitivity of the Q_II_ peak to pressure differed between strains: in *pem2*Δ it was detectable up to 2000 bar, while in WT and *psd1*Δ + CHO at 1 bar, it dissipated by 500 bar. The midpoints of barotropic phase transitions were used to construct phase diagrams (Fig. 2B). In parallel with phase analysis, we measured lipid packing in the same samples across a P/T grid using high-pressure fluorescence spectroscopy (HPFS). Liposomes were stained with C-Laurdan, a solvatochromic dye whose fluorescence correlates with lipid packing. While we observed some differences in C-Laurdan Generalized Polarization (GP) values between the strains (Fig. 2C), they were modest and did not correspond to changes in L_α_ → L_β_ transition temperatures (Fig. 2C, Supp. Fig. 2).

Our biophysical analyses showed that changes in PE vs. PC headgroup metabolism primarily affected curvature-driven phase behavior, and not membrane fluidity, thus allowing us to ask whether lipid curvature dictates pressure tolerance as previously observed in bacterial cells (23). We assessed the pressure tolerance of each strain by measuring growth during and survival following hydrostatic pressure incubation. For the former, relative pressure effects were calculated via changes in cell density (ΔOD_600_) compared to identical 1 bar incubations (ΔΔOD_600_) (Fig 2D).

Cells lacking Psd1, with strongly reduced PE levels, showed a growth defect through this analysis. Survival, measured by spotting culture aliquots after pressure incubation, was also strongly reduced in *psd1*Δ, as well as in *psd1*Δ + CHO, which exhibited a moderate PE reduction. Thus, loss of PE synthesis, which reduces accessibility to non-lamellar phases, renders cells sensitive to high pressure that further reduces lipid curvature.

### Cells under pressure compensate by increasing lipidome curvature, but not membrane fluidity

While *psd1*Δ cells with reduced access to non-lamellar phases showed a growth defect at pressure compared to WT, we did not observe the corresponding growth enhancement for *pem2*Δ cells that showed increased access to non-lamellar phases. Importantly, our biophysical analyses were carried out in cells grown at 1 bar, so they did not account for any changes that pressure could cause to yeast lipid metabolism during extended incubations. We thus considered whether cells might acclimate their lipidomes to pressure, thereby improving their fitness. Because HPSAXS analysis requires large amounts of biomass, we focused this analysis on WT cells grown in identical, large pressure vessels incubated either at 1 or 250 bar for 16 h, from which extracted polar lipids were used for both HPSAXS and HPFS to measure lipid phase properties membrane fluidity, respectively (Fig. 3A).

**Fig. 3:**
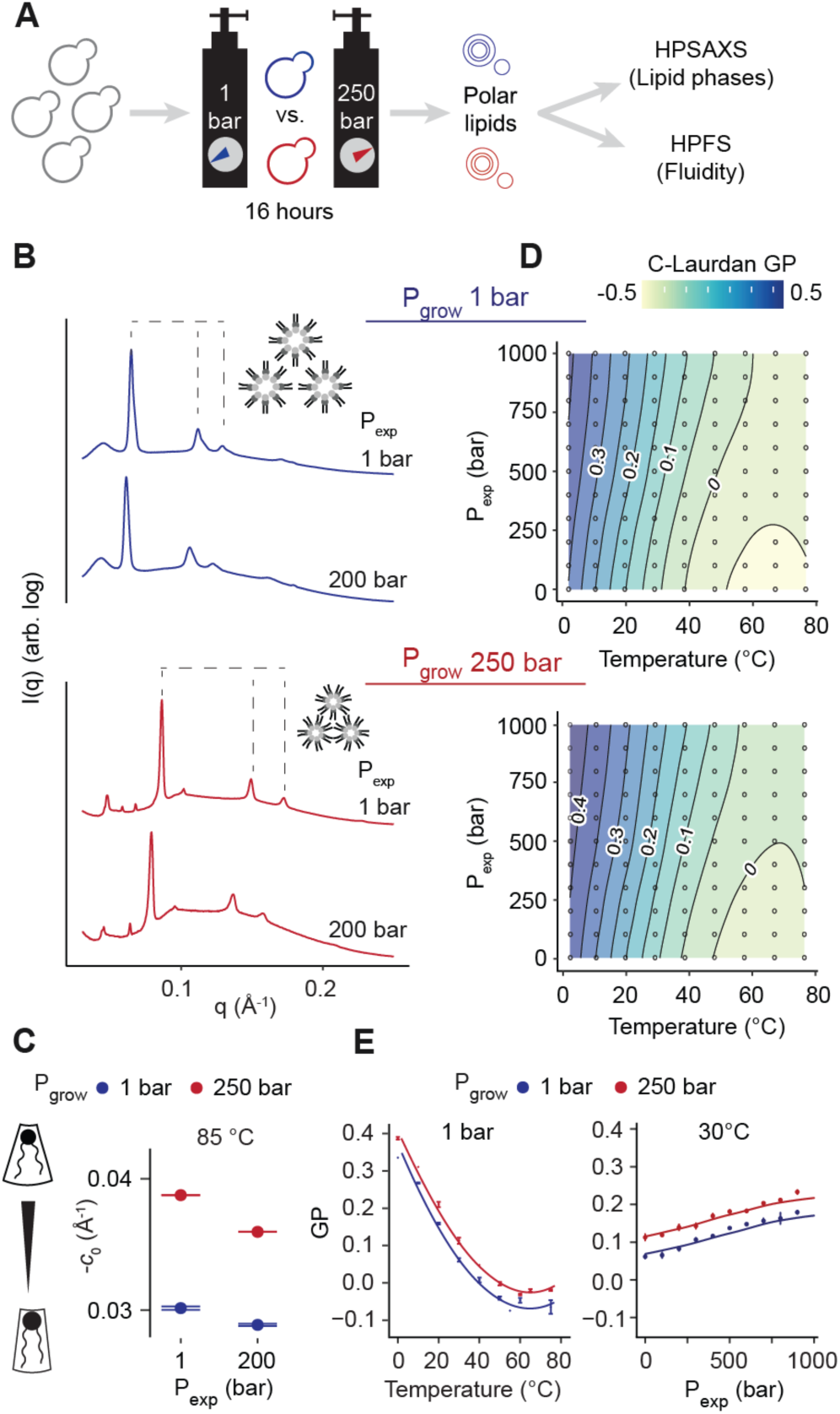
Biophysical signatures of homecurvature acclimation in lipid dispersions from yeast grown under pressure. *(**A**)* Schematic of workflow for high-pressure acclimation experiments. *(**B**)* Representative HPSAXS profiles used to determine the phase of total polar lipid dispersions from W303 yeast grown at 1 and 250 bar pressure. Profiles are shown for extracts at 1 and 200 bar SAXS cell pressure conditions at 85°C for dispersions from yeast grown at 1 and 250 bar, with baselines offset for clarity. Characteristic HII phase peak spacing (1, √3, 2) are labelled in each 1 bar profile along with cartoon insets showing the corresponding difference in Hii size. *(**C**)* Differences in curvature extracted from the SAXS HII phase of yeast grown at 1 and 250 bar pressure measured at 85°C. Lipidome adjustment more than offsets the effect of pressure on curvature, as measured in reconstituted polar lipid liposomes. Error bars indicate SEM from a linear regression fit of curvatures determined at pressures from 1-400 bar. *(**D**)* C-laurdan GP values of liposomes from the same samples measured across a P-T grid with HPFS. Lower GP values correspond to more fluid membranes. Non-linear behavior at higher temperatures corresponds to the non-lamellar phase transition, which occurs at a higher pressure for the extract grown at 250 bar. The color key at the top indicates GP values. *(**E**)* C-Laurdan GP across a single reference pressure (1 bar, left) and temperature (30°C, right). Liposomes prepared from high-pressure extracts show a higher GP value, reflecting reduced membrane fluidity.

In HPSAXS experiments, reconstituted extracts formed non-lamellar phases at elevated temperatures 85°C (Fig. 3B): 1 bar-grown extracts showed a mixture of H_II_ and lamellar signatures, while the 250 bar-grown extracts showed a predominantly nonlamellar mixture of Q_II_ and H_II_ signatures. Taking advantage of the presence of a clear H_II_ phase in both samples, we were able to extract curvature values which correspond to differences in lipid composition in each extract (Fig. 3C). The 250 bar-grown H_II_ phase displayed larger peak spacing corresponding to a curvature *c* of -0.038 +/- 0.00025 Å^-1^ compared to *c* = -0.030 +/- 0.00014 Å^-1^ in the 1 bar-grown sample. *c* here is expected to be slightly more negative than intrinsic curvature *c*_0_ due to chain stretching stress in the H_II_ lattice (29). In parallel, we tested whether pressure elicits a fluidity response in the same samples by measuring C-Laurdan GP across 10-80°C and 1-1000 bar (Fig. 3D). These experiments showed that the 250 bar extracts exhibited reduced fluidity compared to those from cells grown at 1 bar, which is clear at single-temperature or experimental pressure references (Fig. 3E). This difference was in contrast to what would be expected for a homeoviscous acclimation to pressure. Yeast cells thus respond to pressure through a response that maintains lipidome curvature, not membrane fluidity.

### Yeast promote negative curvature by increasing the abundance of the non-bilayer lipid PI

We hypothesized that the propensity of high-pressure extracts to form nonlamellar topologies could be explained by upregulation of high-curvature lipids. We anticipated increased levels in PE and a reduction of PC as the canonical non-bilayer and bilayer-forming phospholipids, respectively. While the PE/PC ratio (0.31 +/- 0.05) did increase in cells grown at 250 bar (0.39 +/- 0.05), there was an overall decrease in both PC and PE (Fig. 4A). Instead, levels of PI increased close to 3-fold compared to cells grown at atmospheric pressure. We also observed changes in the acyl chains across all phospholipids. High-pressure growth caused an increase in monounsaturated phospholipids at the expense of saturated and diunsaturated species (Fig. 4B). There was also an increase in long-chain phospholipids, a pattern previously observed in pressure-adapted ctenophore lipids (26). Lengthening of lipid chains without corresponding unsaturation promotes negative curvature while reducing membrane fluidity; it is thus clearly a curvature-specific response (30).

**Fig. 4:**
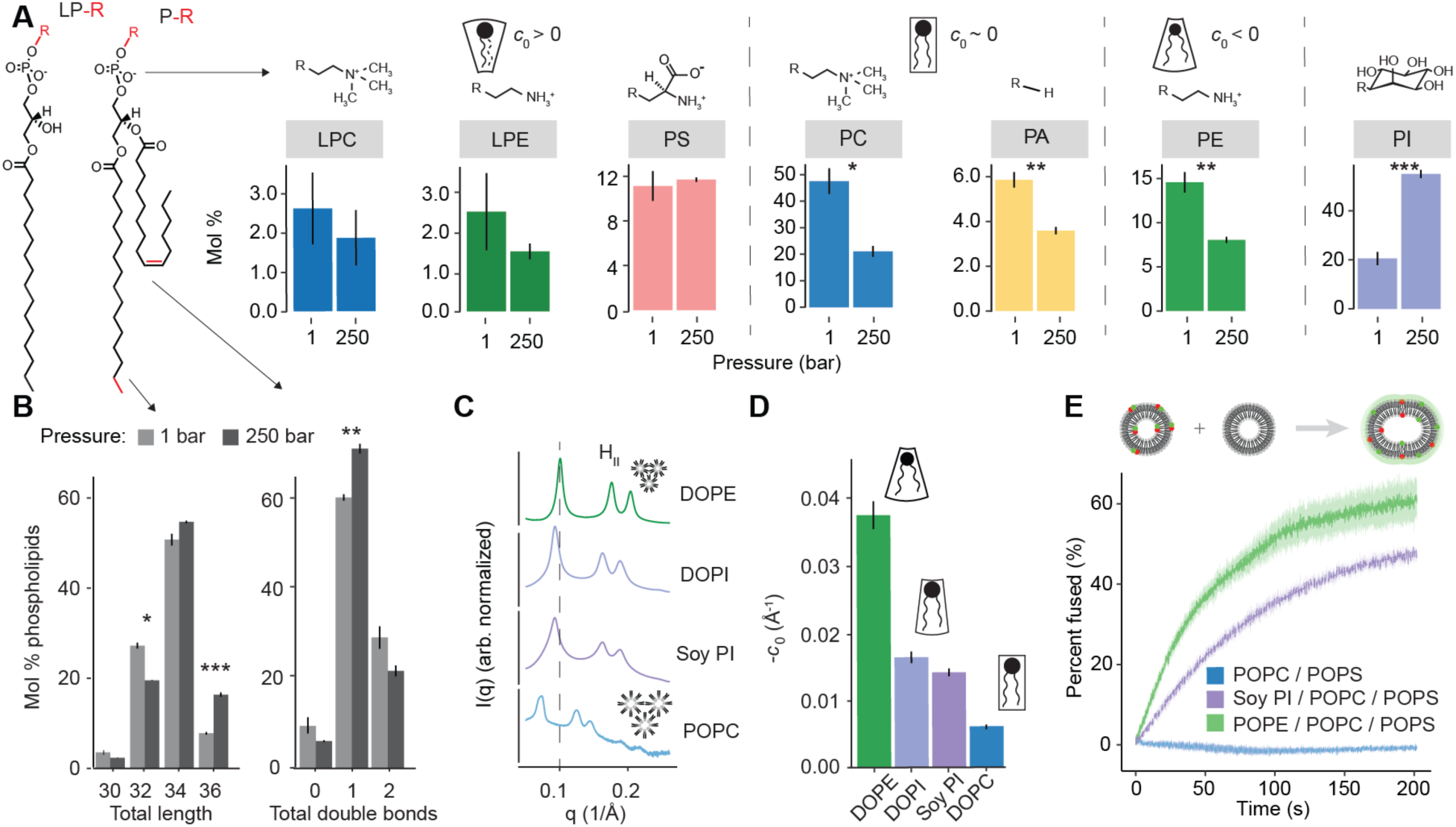
Identification of PI as a curvature-promoting phospholipid that accumulates in response to high pressure. *(**A**)* Abundances of phospholipid classes in yeast grown at 1 or 250 bar pressure, with chemical structures of the representative headgroups above each. Error bars indicate SEM from *N* = 3 independent cultures. * *P* < 0.05, ** *P* < 0.01, *** *P* < 0.001 unpaired two-tailed t-test. *(**B**)* Corresponding total length (left, summed over both chains) and number of double bonds (right) summed over both chains of all phospholipids. Error bars indicate SEM from *N* = 3 independent cultures. * *P* < 0.05, ** *P* < 0.01, *** *P* < 0.001 unpaired two-tailed t-test of total length. (***C***) Representative SAXS profiles of the HII phase of 100% DOPE and hosted DOPI (10 mol %), Soy PI (10%), and POPC (10%) samples in DOPE (90%), all relaxed with 12% (w/w) tricosene at 1 bar, 35 °C. The shift in spacing of the first peak between each lipid is denoted. Leftward shifts represent a reduction in negative curvature. (***D***) Intrinsic curvature (c0) values determined for DOPE, DOPI, Soy PI, and DOPC at 1 bar, 35 °C. Error bars indicate SEM from a linear regression fit of curvatures determined at pressures from 1-1000 bar. (***E***) Lipid mixing assay comparing the fusion efficiency of liposomes with 33:33:33 mol % POPE/POPC/POPS, 33:33:33 mol % Soy PI/POPC/POPS, or 66:33 mol % POPC/POPS. FRET dequenching occurs when one population of vesicles, containing 2 mol % Rh-PE and 2 mol % NBD-PE, fuses with an unlabeled population of vesicles following the addition of 10 mM CaCl2. Intensities are normalized to the assay performed with fully fused vesicles. Only liposomes containing PE or PI as non-bialyer lipids are capable of fusing.

The dominance of PI in pressure-treated yeast cells motivated further investigation of its biophysical properties. Although PI is a ubiquitous eukaryotic phospholipid and is especially abundant in yeast, its phase and curvature properties have been less well characterized than those of its phosphorylated derivatives (31). However, one previous study observed that PI promoted cubic phases in mixtures with PC (32). We first tested the phase properties of Soy PI, which features a mixture of acyl chains but is available in bulk. Soy PI in water (50% w/v) displayed a complex mixture of cubic and lamellar signatures that were stable from 10°C and 80°C and from 1-1000 bar pressure (Supp. Fig. 3A). C-Laurdan measurements reflected an intermediate fluidity between PC and PE, but were more challenging to interpret due to the phase coexistence (Supp. Fig. 3B). To measure PI’s curvature directly, we extrapolated *c_0_* from hosted mixtures of 10 mol % PI in DOPE H_II_-phase tubules (Fig. 4C). For these, the H_II_ phase was relaxed with the hydrocarbon (9Z)-tricosene. Both Soy PI and di-oleoly PI (DOPI), with defined 18:1 chains in both positions, featured similar *c_0_* values of -0.015 Å^-1^ at 1 bar and 35°C (Fig. 4D). This value was between DOPE’s larger negative curvature and DOPC’s near-zero curvature under these conditions. PI in water thus exhibits a negative intrinsic curvature that is intermediate to the other main yeast phospholipids PC and PE.

Negative-curvature lipids facilitate membrane fusion by stabilizing nonbilayer intermediates. To test whether the negative curvature of PI could permit membrane fusion, we performed a FRET-based, Ca^2+^-induced lipid mixing assay of liposomes containing phosphatidylserine (PS), PC, and either PE or PI as the nonbilayer lipid (Fig. 4E). Unlike control liposomes with only PC and PS, those including either Soy PI or PE were fusogenic. Fusion rates with Soy PI were lower than those with PE, consistent with its more moderate negative curvature.

### Multiple mechanisms of homeocurvature acclimation across yeast and mammalian cells

While we noted an increase in PI levels, pressure induced multiple lipidome changes in yeast cells, affecting both the distribution of phospholipid head groups (Fig. 4B), chains (Supp. Fig. 4), and abundance of lysolipids, which contribute positive intrinsic curvature (33). To globally account for these changes, we extended a mathematical model (c0operate) for estimating a phospholipid curvature index (PLCI) that assumes ideal mixing between phospholipid species and accounts for differences in their head groups and acyl chains (23, 34). Analyzing the *S. cerevisiae* data (Fig. 5A), PLCI becomes more negative (higher curvature elastic stress) in 250 bar-grown cells and this change is primarily driven by substitution of PI for PC (Fig. 5B). We then performed a similar analysis on a different budding yeast, *Lachancea kluyveri*, that diverged from *S. cerevisae* at least 100 million years ago (35). *L. kluyveri* also responded to pressure by increasing PI levels, alongside a notable reduction in positively-curved PS. Acyl chains of pressure-grown *L. kluyveri* increased in length and unsaturation, employing its ability to produce polyunsaturated fatty acids that *S. cerevisiae* does not (Supp. Fig. 4). Overall, both yeasts displayed a similar response of PLCI to 250 bar (Fig. 5B).

**Fig. 5:**
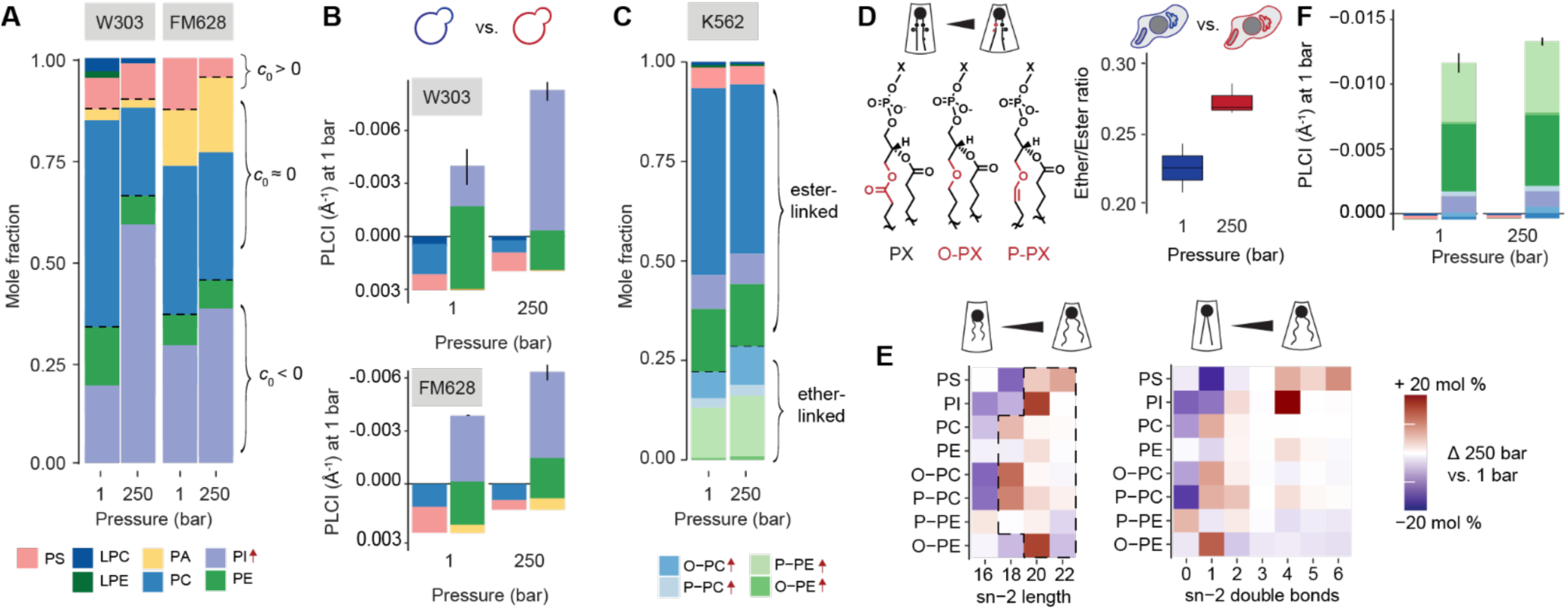
Lipidome curvature acclimation across cell types. (***A***) Phospholipid abundances for *S. cerevisiae* (W303) and *L. kluyveri* (FM628) lipidomes from cultures grown at 1 and 250 bar pressure. Values represent the mean phospholipid molar fraction of *N* = 3 independently grown cultures. Arrows next to the key indicate classes with increased abundance. (***B***) PLCI of W303 and FM628 lipidomes during growth at 1 or 250 bar. The y-axis is inverted to emphasize increases in negative *c0*. Individual lipid contributions to positive *c0* are plotted to the left, and those contributing to negative curvature on the right. The latter are offset by the contribution of the former to show the change in net PLCI. Error bars indicate SEM, *N* = 3. *P* = 0.013 (W303, top), *P* = 0.027 (FM628, bottom); unpaired one-tailed t-test. (***C***) Phospholipid classes of K562 suspension cells incubated at 1 or 250 bar for 6 h. Ether-linked lipids (P-PE, O-PE, O-PC, and P-PC) increase in abundance. (***D***) Structures of ester (PX), alkyl ether (O-PX), and vinyl ether (P-PX) phospholipids, whose total abundance compared to ester-linked lipids increases with pressure (right). Error bars indicate SEM from *N* = 3 independent cultures. *P* = 0.047, unpaired two-tailed t-test. (***E***) Heatmaps showing increases (red) in phospholipids with longer (right) and more unsaturated (left) sn-2 chains in cells grown at 250 bar vs. the 1 bar control. These correspond to decreases in shorter and more saturated chains (blue). The dashed box highlights longer-chain species that promote negative curvature. (***F***) PLCI analysis for K562 shows higher negative lipidome curvature at 250 bar. Error bars indicate SEM from *N* = 3 independent cultures. *P* = 0.054, unpaired one-tailed t-test. Values are plotted as in panel B.

Though PI is abundant in yeast, it is a smaller component in mammalian cells and is absent in most bacteria. We thus asked whether curvature maintenance is a general biophysical phenomenon in these systems or is limited to regulation of PI levels. We subjected K562, a human leukemia suspension cell line, to incubations at 1 or 250 bar in the same chambers used in the yeast experiments, albeit for shorter 6 h incubations. While pressure-induced toxicity prevented longer treatments, we did observe lipidome remodeling due to pressure (Fig. 5C). Notably, we observed an increase in ether phospholipids, which include P-PE, monoalkyl PE (O-PE), plasmenyl PC (P-PC), and monoalkyl PC (O-PC), and a corresponding reduction in PC (Fig. 5D). Ether linkages contribute directly to negative curvature and are an adaptive feature of ctenophores specialized for high-pressure (23, 30). K562 cells grown at 250 bar also showed an increase in phospholipid chain length, a signature of a curvature response (30), and unsaturation, which is shared by both curvature and fluidity responses (Fig. 5E). These changes were concentrated in the sn-2 chain position and were observed in almost all phospholipid classes (Supp. Fig. 5). Globally, when accounting for these changes, the PLCI became more negative in pressure-treated K562 cells (Fig. 5F) – a similar effect as in yeast but through different lipidic levers. The extent of curvature acclimation in K562 could be underestimated through these calculations due to a paucity of reference *c_0_* values for phospholipids containing long-chain PUFAs.

As a model bacterium, *Escherichia coli* cannot produce any of the above curvature-associated lipid classes (PI, ether PE and PCs) but has been previously shown to maintain access to non-lamellar phases in response to temperature (19). However, when grown at even 500 bar, we failed to detect any changes to mean phospholipidome curvature in *E. coli* BL21 cells (Supp. Fig. 6A). The ratio between PE and PG headgroups remained constant, as did the number of double bonds in acyl chains (Supp. Fig. 6B). While there were some small changes in the distribution of chains, longer or more unsaturated chains did not accumulate at high pressure (Supp. Fig. 6C). These results suggest that curvature responses in *E. coli* are likely a side-effect of thermal acclimation. They are also consistent with work showing that fatty acid unsaturation is regulated by temperature sensitivities of enzymes in *E. coli*’s type II fatty acid synthesis (12), and so would not respond directly to loss of curvature frustration induced by high-pressure incubations.

## Discussion

Non-bilayer lipids must be regulated to maintain cell membranes that are stable yet support dynamic functions dependent on curvature elastic stress. Previous studies found that non-bilayer lipid classes are regulated across the cell cycle (2, 22, 36) and in response to perturbations in lipid metabolism (20). However, it has not been known whether this regulation is in response to curvature stress itself – if cells directly sense the shape of their lipids – or occurs as a side effect of fluidity homeostasis. To address this question, we employed hydrostatic pressure to directly perturb lipid curvature. The eukaryotic cells we tested respond to high pressure by increasing curvature-promoting chemical features across the phospholipidome, even at the expense of membrane fluidity.

Acclimation of phospholipids to pressure showed a tendency to overshoot with respect to curvature: the estimated compensatory change of PLCI in the three systems analyzed are 1.5 (K562) to 3 (*S. cerevisiae* W303) fold higher than the increase in the *c_0_* of model lipids like DOPE at 250 bar. Such an overcompensation was previously observed in native lipidome adaptation across deep-sea ctenophores (23) and transiently during cold acclimation in bacteria (12). Overshoot could result from a need for the phospholipidome to compensate for other abundant membrane components – transmembrane proteins and lipid classes like sphingolipids – that inhibit non-lamellar topologies and are not amenable to acclimation. Our analyses also did not account for subcellular compositions or asymmetric distributions across bilayer leaflets, like in the plasma membrane where non-bilayer lipids like PE are found on the inner leaflet. It is likely that maintenance of curvature stress at specific sites within cells or membrane leaflets drives global lipidome response. Nonetheless, whole cell lipidomes yield general acclimatory trends that are likely to be relevant across different compartments, even if they do not reflect the curvature stress at any one site.

The systems analyzed here show a range of lipidome responses that underlie curvature regulation. In yeast, high pressure increased the PE/PC ratio, but more dramatically increased levels of PI. PI was previously observed to promote the Q_II_ phase in PC-hosted experimental systems (32) but its curvature had not been previously determined, so there has been disagreement over its classification as a non-bilayer lipid (37, 38). We find that PI features a negative curvature when measured in a DOPE-hosted H_II_-phase system and forms a mixture of lamellar and non-lamellar phases in isolation. Consistent with this property, substitution of PI for PE fulfills the requirement of a nonbilayer lipid in permitting membrane fusion. The anionic nature of PI and its bulky headgroup would argue against a negative intrinsic curvature. However, NMR and neutron diffraction measurements show that PI’s inositol takes on extended conformations from the bilayer plane that maximize intra- and inter-molecular hydrogen bonding and would reduce bulk proximal to the bilayer (39–42). The inositol ring of PI can undergo hydrogen bonding at five positions; hydrogen bonding between lipids contributes to negative curvature (43) and is a general feature on sugar-containing lipids that form non-lamellar phases (44–46). Although PI has only a moderate negative curvature, it does not show the high ordering and melting temperature of PE (47), suggesting a rationale for why its abundance is modulated more widely by yeast. Consistent with this model, we had previously observed that PI substitutes for PE in yeast with highly saturated fatty acid pools (48).

In mammalian cells, PUFA chains and plasmenyl linkages help to mitigate the high T_m_ of PE, potentially allowing it to serve as the primary curvature modulator. K562 cells respond to the reduction in lipid curvature at high pressure by increasing ether-linked phospholipids and through remodeling of the sn-2 acyl chains. Sn-1 -alkyl and -plasmenyl linkages on PC and PE, which yeast are not capable of synthesizing, are structural features that likely contribute directly to negative curvature and promote curvature-dependent properties like membrane fusion (8) or invaginations of the plasma membrane (49). Ether lipid metabolism could thus act as a regulator of membrane curvature stress in mammalian cells, a role that is relevant to its dysregulation in numerous metabolic disorders and neurodegenerative diseases (50, 51). Both yeast and K562 also show changes in phospholipid chains, but this effect is larger in the latter given its larger repertoire of fatty acids and their incorporation by the Lands cycle. Longer phospholipid chains represent a curvature-specific lipidome response because they promote non-lamellar phases, but also act against membrane fluidity and promote membrane gelling (30).

A question introduced by this work concerns the mechanisms by which cells could sense membrane curvature stress or related properties to maintain homeostasis. Because the cells analyzed are not derived from marine systems, these mechanisms are not likely to sense hydrostatic pressure directly. Instead, we expect membrane-associated sensors to control multiple aspects of phospholipid metabolism through direct or transcription-mediated pathways.

Choline cytidyl transferase (CCT), which catalyzes the committed step in PC synthesis, is dependent on lipid curvature. CCT can be inhibited by non-bilayer lipids in liposomes, suggesting that it is regulated by curvature stress (52, 53). This phenomenon can explain the reduction in PC synthesis we observe in pressure-treated cells. Low PC synthesis derepresses expression of *INO1* (54), which encodes for inositol-3-phosphate synthase (Ino1) that catalyzes the first step of PS synthesis. Earlier microarray studies showed a stimulation of *INO1* expression in pressure-treated yeast cells (55), consistent with this model for its regulation. Other mechanisms for the curvature regulation observed here are less clear. The Lands cycle generates tissue-specific acyl chain compositions in mammals (56), but few studies have investigated how its enzymes are regulated by the membrane’s physical state. Similarly, regulation of ether lipid metabolism -– a complex pathway occurring across both the peroxisome and the ER – has yet to be elucidated.

Motivated by homeoviscous adaptation, acclimation to maintain access to non-lamellar phases was previously proposed through measurement of the L_α_ to H_II_ transition in bacterial lipids extracted from different growth temperatures (19). Even though synthetic manipulation of lipidome curvature modulates *E. coli* pressure fitness (19), our data suggests that its natural phospholipid and fatty acid metabolisms are unlikely to be regulated by lipid curvature itself. Given that the pressure-regulated lipid classes we observed are more typical of eukaryote lipidomes, it is possible that curvature-based regulation is ubiquitous in eukaryotes but found only in pressure-adapted bacteria (30, 57). Eukaryotic cells feature intracellular trafficking networks that depend on the ability of membranes to undergo fission and fusion, two processes strongly dependent on lipid curvature, as well as more complex membrane protein machinery. Homeocurvature acclimation could have thus arisen in cells that require fine tuning of membrane dynamics for homeostasis of their organelles or other yet undefined processes. Hydrostatic pressure, as a physical modulator of lipid curvature, represents a powerful experimental tool to identify these functions.

## Acknowledgements

Alexander Sodt and Edward Lyman contributed helpful discussions. Kailash Venkatraman provided experimental support and comments. Richard Gililan and Qingqiu Huang assisted in beamline measurements. Research was funded by the National Science Foundation (NSF) (MCB-2046303 and MCB-2316458 to I.B.), the Office of Naval Research (N00014-23-1-2543 to I.B.), the National Institutes of Health (NIH) (5T32EB009380-14 to D.M., GM139641 to E.A.D., GM142960 to I.B.), and NASA (postdoctoral fellowship 0017-NPP-MAR22-A-Astrobio to J.R.W.). SAXS measurements were performed at the Center for High-Energy X-ray Sciences (CHEXS), which is supported by the NSF award DMR-2342336, and the MacCHESS resource is supported by NIH award 1-P30-GM124166.

## Materials and Methods

### Yeast strains and growth conditions

All *S. cerevisiae* strains used in this study are described in Supplementary Table 1. Cells were grown in complete supplement mixture (CSM) (0.5% Ammonium Sulfate, 0.17% yeast nitrogen base without amino acids and 2% glucose) lacking amino acids as appropriate for selection. Yeast mutants were generated by PCR-based homologous recombination, ORFs *(PSD1, PEM2)* were replaced by *HIS3*. For standard incubations, cells were grown overnight at 30°C in a temperature-controlled shaker (200 RPM) in yeast extract-peptone-dextrose (YPD) medium using 14 mL culture tubes (18 mm diameter) with a snap/vent cap (Greiner). Cells were then back-diluted into 5 mL fresh CSM and grown until late-exponential phase (OD 1-1.5) prior to analysis.

For high-pressure cell fitness and lipidomics analyses (Figures 2, 4, and 5A-B), cultures were pre-incubated in YPD in a shaker overnight at 30°C. Samples were then back-diluted to OD 0.1 in CSM containing 2% glucose and loaded into sterile polyethylene 4.5 mL transfer pipettes (Thermo Scientific), the bulbs of which were subsequently heat-sealed with a manual impulse sealer (YXQ)(58). The bulbs containing cultures were then loaded into each of two stainless steel pressure vessels (59) filled with distilled water and pre-warmed to 30°C. A manual rotary pump (High Pressure Equipment Co.) was used to bring one vessel to 250 bar, and a second was left at ambient pressure as a 1 bar control. All pressure vessels were clamped horizontally to an orbital shaker and shaken at 250 RPM overnight at 30°C. Following high-pressure incubation, vessels were decompressed over the course of ∼30s. OD_600_ was measured for all replicate cultures of each strain at each pressure and a qualitative survival screen was performed as follows. Each culture was serially diluted fivefold, yielding aliquots that were 5^0^- to 5^5^-fold dilutions of the decompressed culture. Six sets of these were transferred to a YPD agar plate using a 48-pin tool (V&P Scientific) and each plate was incubated at 30°C for approximately 24 h, then photographed. The remainder of each decompressed culture was pelleted at 3000 RCF, 5 min at 25°C and frozen at -80°C. Total lipids were later extracted from the frozen pellets by the Bligh-Dyer method (60) and samples incubated at 1 and 250 bar were analyzed as described below.

For high-pressure biophysical analyses (Figure 3), single colony cultures were pre-incubated in YPD in a shaker overnight at 30°C. Samples were then back-diluted 1:100 into 2.5 L YPD in bleach-sterilized bags made from Mylar™ tube stock containing a magnetic stir bar. The bags were heat-sealed with a manual impulse sealer, leaving zero headspace. The cultures were loaded into custom-built 4 L titanium pressure vessels filled with distilled water and prewarmed to 30°C using water jackets. One of the pressure vessels was brought to 250 bar hydrostatic pressure using a manual hydraulic hand pump (Enerpac). Cultures were incubated overnight at 30°C with stirring at ∼200 RPM. Following high-pressure incubation, the 250-bar vessel was decompressed and the bag contents of the culture bag were pelleted at 3000 RCF and 25°C. Pellets were flash frozen, total lipids were extracted by the Bligh-Dyer method and samples incubated at 1 and 250 bar were analyzed as described below.

### Mammalian cell culture and pressure incubation

K562 cells (ATCC) were grown and maintained in Iscove’s Modified Dulbecco’s Medium (IMDM; Gibco) supplemented with 1% penicillin-streptomycin (Gibco) and 10% fetal bovine serum (Gibco). For pressure incubation, N=3 sterile transfer pipette bulbs were filled with 4 mL of K562 cells at ∼7.5 x 10^6^ cells/mL. After eliminating air bubbles, the bulbs were heat-sealed as described above and loaded into pressure vessels filled with water and pre-warmed to 37°C. A manual rotary pump (High Pressure Equipment Co.) was used to pressurize one vessel to 250 bar; the other was left at ambient pressure as a 1 bar control. Both pressure vessels were incubated at 37°C for 6 h, after which the 250-bar vessel was decompressed over ∼30 s. OD_600_ was measured for all three replicates at each pressure. Each decompressed culture was pelleted at 3000 RCF for 5 min at 25°C and frozen at -80°C. Total lipids were later extracted from the frozen pellets and analyzed as described below.

### Bacterial cell culture and pressure incubation

Bacterial lipidome data were repurposed from a previously published experiment for this study (23). Starter cultures of BL21 (DE3) *E. coli* containing pET28 (empty control) plasmids were grown in LB + 50 mg/L kanamycin (Kn) while shaking at 250 RPM, overnight at 37°C. Each starter was diluted 1:100 v:v into LB + 100 mM HEPES pH 7.4 + Kn and incubated micro-aerobically while shaking at 250 RPM for 24 h at 37°C. Starter tubes were connected to a water airlock and the headspaces flushed with N_2_ prior to culturing. At OD600 = 0.2, IPTG was added to 0.1 mM final concentration, headspaces were flushed again, and microaerobic culture was continued for another 24 h. Following this period, all cultures were normalized to OD_600_ = 0.2 using the same medium and 3 mL aliquots were loaded into 5 mL polypropylene syringes each containing three 3 mm glass beads for agitation. Gas was manually purged from the syringes and each was sealed with a Luer-lock cap. Three syringes containing each strain were loaded into pressure vessels filled with distilled water and pre-warmed to 37°C. A manual rotary pump (High Pressure Equipment Co.) was used to pressurize one vessel to 500 bar and the other was left at ambient pressure as a 1 bar control. The pressure vessels were clamped horizontally to an orbital shaker and shaken at 250 RPM for 48 h at 37°C. Following high-pressure incubation, vessels were decompressed over the course of ∼30 s. OD_600_ was measured for all three replicate cultures each pressure. Each decompressed culture was pelleted at 3000 RCF for 5 min at 25°C and frozen at -80°C. Total lipids were later extracted from the frozen pellets by the Bligh-Dyer method and samples incubated at 1 and 500 bar were analyzed as described below.

### Lipidomic analysis

Mass spectrometry-based analysis of yeast phospholipids (Figures 2, 4, 5A-B; Supp. Fig. 4) was performed by Lipotype GmbH (Dresden, Germany) as described (61). Lipids were extracted using a chloroform/methanol procedure (62). Samples were spiked with internal lipid standard mixture containing: diacylglycerol 17:0/17:0 (DAG), ergosterol ester 13:0 (EE), phosphatidate 17:0/14:1 (PA), phosphatidylcholine 17:0/14:1 (PC), phosphatidylethanolamine 17:0/14:1 (PE), phosphatidylglycerol 17:0/14:1 (PG), phosphatidylinositol 17:0/14:1 (PI), phosphatidylserine 17:0/14:1 (PS) and triacylglycerol 17:0/17:0/17:0 (TAG). After extraction, the organic phase was transferred to an infusion plate and dried in a speed vacuum concentrator. The dry extract was re-suspended in 7.5 mM ammonium formate in chloroform/methanol/propanol (1:2:4; v:v:v). All liquid handling steps were performed using the Hamilton Robotics STARlet robotic platform with the Anti Droplet Control feature for organic solvent pipetting. Samples were analyzed by direct infusion on a Q Exactive mass spectrometer (Thermo Scientific) equipped with a TriVersa NanoMate ion source (Advion Biosciences). Samples were analyzed in both positive and negative ion modes with a resolution of R_m/z=200_=280,000 for MS^1^ and R_m/z=200_=17,500 for MS^2^ experiments, in a single acquisition. MS^2^ was triggered by an inclusion list encompassing corresponding MS mass ranges scanned in 1 Da increments (63). Both MS^1^ and MS^2^ data were combined to monitor EE, DAG and TAG ions as ammonium adducts; PC as formate adducts and PA, PE, PG, PI and PS as deprotonated anions. Data were analyzed using LipotypeXplorer, a proprietary software developed by Lipotype GmbH, which is based on LipidXplorer (64, 65). Data post-processing and normalization were performed using an in-house developed data management system. Only lipid identifications with a signal-to-noise ratio >5, and a signal intensity 5-fold higher than in corresponding blank samples were considered for further data analysis.

Mass spectrometry-based analysis of BL21 and K562 phospholipids (Figures 5C-E, Fig. 5, 6) was performed as previously described (23). Lipid extracts were brought to dryness and reconstituted in 18:1:1 v:v:v IPrOH:CH_2_Cl_2_:MeOH containing deuterated internal standard (Avanti) for each of the 11 lipid classes analyzed, which include the major classes of phospholipids. These allow for semi-quantification of species within these classes. Other potentially present lipid species, such as ceramide derivatives, glycosylated lipids, and non-sterol terpenes, were not quantified using this method. Polar lipids were measured according to a previously described method (66), with slight modifications. The Vanquish UHPLC (Thermo Fisher Scientific) was interfaced with a Q Exactive mass spectrometer (Thermo Fisher Scientific). A Waters T3 1.6 µM 2.1 mm x 150 mm column was used for chromatographic separation using a step gradient from 25% buffer A (10 mM ammonium formate and 0.1% formic acid in water) to 100% buffer B (70/30 v/v isopropanol/acetonitrile with 10 mM ammonium formate and 0.1% formic acid) over 35 min. Flow rate was set at 0.3 mL/min. Lipid samples were analyzed using data-dependent acquisition (DDA) with an NCE of 30 in negative mode and an NCE of 25 in positive mode. Ion source parameters were: Sheath Gas 48 AU; Aux Gas 11 AU; Sweet Gas 1 AU; Spray Voltage 3.5 kV for positive mode and 2.5 kV for negative mode; Capillary Temp. 250°C; S-lens RF level 60 AU; Aux Gas Heater Temp. 413°C. All ions in the mass range of 200-1200 m/z were monitored. MS^1^ resolution was set at R_m/z=200_=70,000 with an automatic gain control (AGC) target of 1e6 and Maximum IT of 200 ms. For MS^2^, R_m/z=200_=17,500 with an AGC target of 5e4, fixed first mass of 80 m/z, and Maximum IT of 50 ms. The isolation width was set at 1.2; Dynamic Exclusion was set at 3 s. Lipids were identified and quantified with Lipid Data Analyzer Software (67). All data were analyzed from normalized intensities relative to exactly measured internal deuterated standards (SPLASH LIPIDOMIX, Avanti Polar Lipids).

Due to identical fragmentation patterns, O-PE and P-PE species were manually separated by retention time after Lipid Data Analyzer identification of P-PE species based on MS^2^ fragmentation products: PE head group (m/z = 196.038), carboxy fragment (m/z dependent on chain length), and alkenyl sn-1 chain fragment (m/z = 239.235, 265.253; dependent on chain length). It was possible to distinguish O-PE based on retention time following the principle that alkyl ether species elute sooner than their corresponding vinyl ether counterparts, putatively due to differences in H-bonding imparted by the cis-alkene bond.

To calculate PLCI from lipidomic data, c0operate scripts (68) were used based on inputted lipidomic data and literature values of phospholipid intrinsic curvatures (34) as previously described (23, 30).

### SAXS sample preparation

Total lipid extracts were separated into polar lipids and neutral fractions on 500 mg prepacked silica columns (Thermo Scientific) using a previously validated method (69). Polar lipids eluted in approximately 20 mL CHCl_3_ were dried and weighed in tared 2 mL vials. For SAXS analysis, 5 mg polar lipids were dissolved in 100 µL cyclohexane (C_6_H_12_, Fisher Chemical). 50 µL aliquots were transferred to 1.5 mm ID capillary melting point tubes (Kimble Chase 34505-99), frozen at - 80°C, and lyophilized for 1 h. Samples were rehydrated by the addition of 10 µL vacuum-degassed MilliQ water, then subjected to five rounds of freeze-thaw in an isopropanol-dry ice bath and homogenization with a 20 µL Wiretrol II microdispenser (Drummond Scientific) inside the melting point tube. The bottom of the tube was then scored, cut, and carefully wiped, and the suspension inside transferred to a disposable sample holder (70). This thick suspension (50% w/v lipid) was settled to the bottom of the holder centrifugation (500 RCF, 5 min at 4°C) and topped with another 5 µL degassed water to ensure full hydration. A 22G x 100 mm needle (Air-Tite) was then used to add a pressure-transmitting cap of high vacuum grease (Dow Corning) to the top of the water.

### SAXS data acquisition and analysis

HPSAXS data were collected on beamline 7A at the Cornell High Energy Synchrotron Source using a custom-built temperature-controlled pressure cell (70). Photon energy was 14 keV, spot size was 200 x 250 µm (W x H), and scattering data were collected for 0.005 Å-1 ≤ q ≤ 0.7 Å-1 using an EIGER 4M detector (Dectris) within the beamline vacuum. 1 s exposures were taken at a flux of 1.8-3.6x10^11^ photons/s. Five initial exposures were used to test each sample for radiation sensitivity, then pressure or temperature up- and down-sweeps were made starting at the lowest temperature or pressure and proceeding to the highest. The sample was equilibrated for at least 60 s per 100 bar of pressure change and for 30 min following each 5°C change in temperature. Because photon scattering by the lipid dispersions was strong and background was low (as assessed by shooting a sealed capillary containing the same degassed water used to hydrate the lipids), no background subtraction was used. SAXS images were azimuthally integrated using BIOXTAS RAW software (71).

### C-Laurdan GP determination and analysis

The same polar lipid extracts used for HPSAXS analysis were used to analyze lipid packing via the general polarization (GP) of the C-laurdan solvatochromic probe. HPFS was used to measure general polarization (GP) across a temperature-pressure grid. Polar lipid dispersions were made by drying lipid films under N_2_ and high vacuum for 30 min, then resuspending them in buffer (20 mM HEPES, 2 mM EDTA, pH 7.4) to concentrations of approximately 1 mM lipid. Samples were then subjected to 4X freeze-thaw cycles and subsequently extruded using a hand-held device (Avanti Polar Lipids) incorporating a 200 nm polycarbonate track-etch filter (Whatman) at room temperature. The resultant unilamellar vesicles were stained with 2.5 μM C-Laurdan and tumbled for 30 min at room temperature. Vesicles were then loaded into a quartz cuvette and analyzed in a high-pressure optical cell (ISS Inc.) connected to a 65x SyriXus syringe pressure pump (Teledyne ISCO) and a water bath (NESLAB) for automated pressure and temperature control coordinated by a PC running a Python script (72). The pressure cell was mounted in a Cary Eclipse fluorospectrometer (Agilent Cary). An excitation wavelength of 340 nm was used with the multiwavelength setting acquiring dual emission wavelengths of 440 nm and 490 nm with a 2.5 nm excitation slit and a 5 nm emission slit. Fluorescence data were collected in Kinetics mode concurrent with automated traversal of the temperature-pressure grid. Timestamps were used to co-register temperature and pressure with fluorescence data, which were then analyzed using R scripts.

### Lipid mixing fusion assay

A FRET-based lipid mixing assay was performed to compare fusion rates by fluorescence dequenching of liposomes with equimolar POPC, POPS, and Soy PI or POPE. As a negative control, a condition with only equimolar POPS and POPC was tested. Lipid stocks in chloroform were mixed in equimolar amounts. The chloroform was evaporated to form thin lipid films using N_2_ for 10 min and subsequently kept under high vacuum for ≥1 h to remove residual organic solvent. The lipid film was hydrated with Buffer A (100 mM NaCl, 5 mM Na HEPES, 0.1 mM EDTA, pH 7.4) and tumbled for 1 h. To form unilamellar 200 nm vesicles, the lipid dispersions were subjected to 4 freeze-thaw cycles and extrusion through a 200 nm membrane 21 times. One population of fluorescence-quenched liposomes, containing Rh-DOPE (2 mol %) and NBD-DOPE (2 mol %) (Avanti Polar Lipids), were mixed with a population of liposomes lacking fluorophores. The emission of NBD-PE dequenching was measured following the addition of 10 mM CaCl_2_ using a Cary Eclipse fluorospectrometer set to 463/536 nm ex/em. Fluorescence intensity was recorded in Kinetics mode for 5 min to allow reactions to reach equilibrium. Fluorescence of a fully dequenched control with 1 mol% NBD-DOPE and 1 mM Rh-DOPE was recorded to permit normalization and determination of total % fused vesicles.

## Supporting Information

**Supplementary Figure 1:**
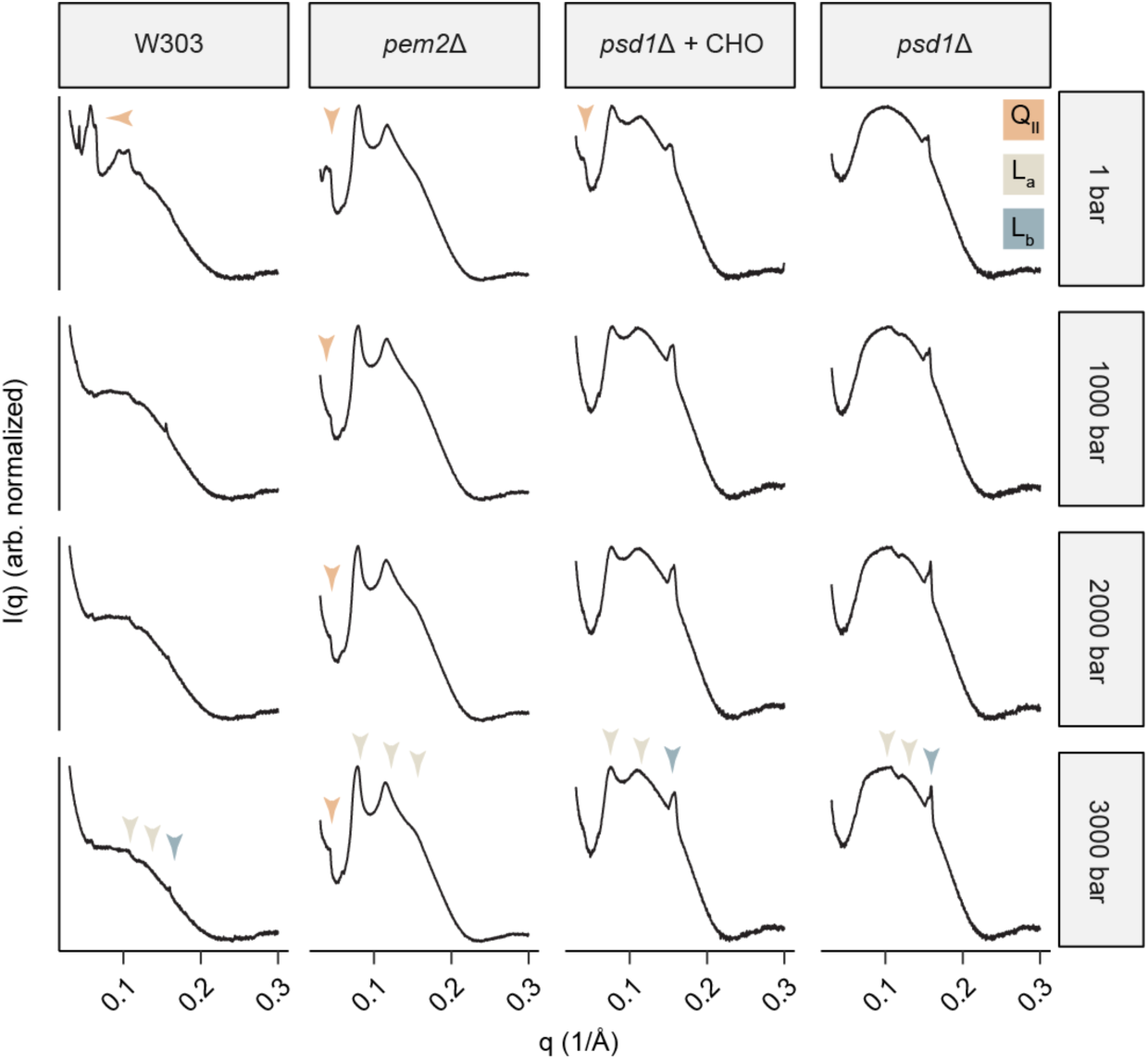
SAXS profiles of polar lipid extracts from yeast strains varying in PE/PC ratio under pressures at 85°C and 1-1000 bar; a subset of the data used to construct Fig. 2B.

**Supplementary Figure 2:**
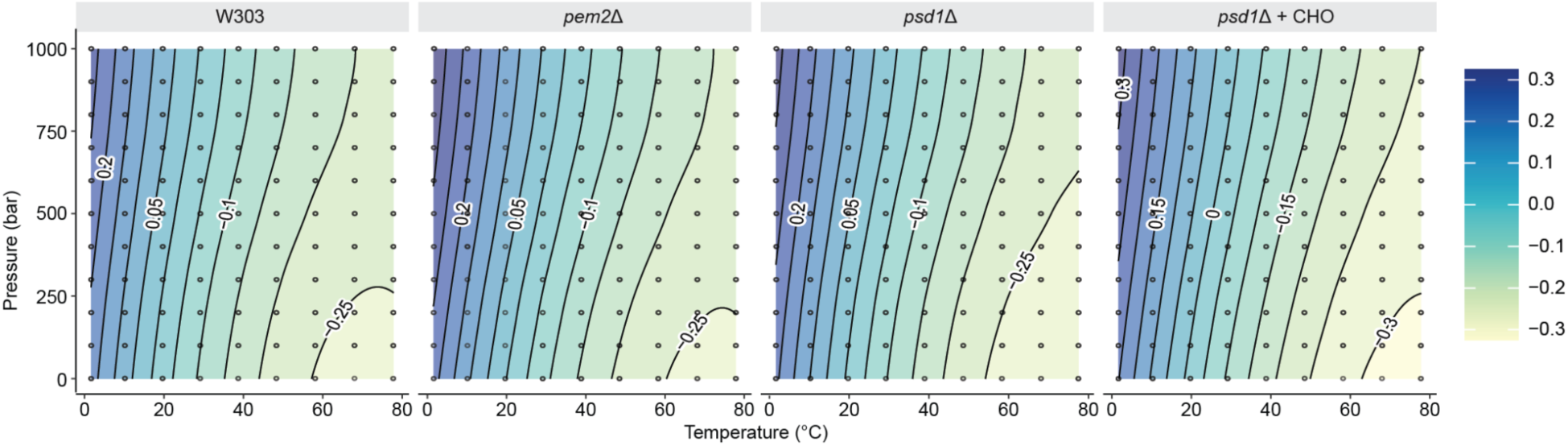
Pressure-temperature phase diagrams of C-Laurdan GP values of polar lipid extracts from PE/PC mutant yeast strains. GP was determined for temperatures and pressures between 0-80°C and 1-1000 bar. Lines show fitted contours of constant GP values across temperature and pressure, while the scale key represents the coding of GP value for color. Lipid extracts show similar fluidity values. The horseshoe trend lines of GP at high temperatures observed in all systems except for *psd1*Δ reflects a non-lamellar phase transition.

**Supplementary Figure 3:**
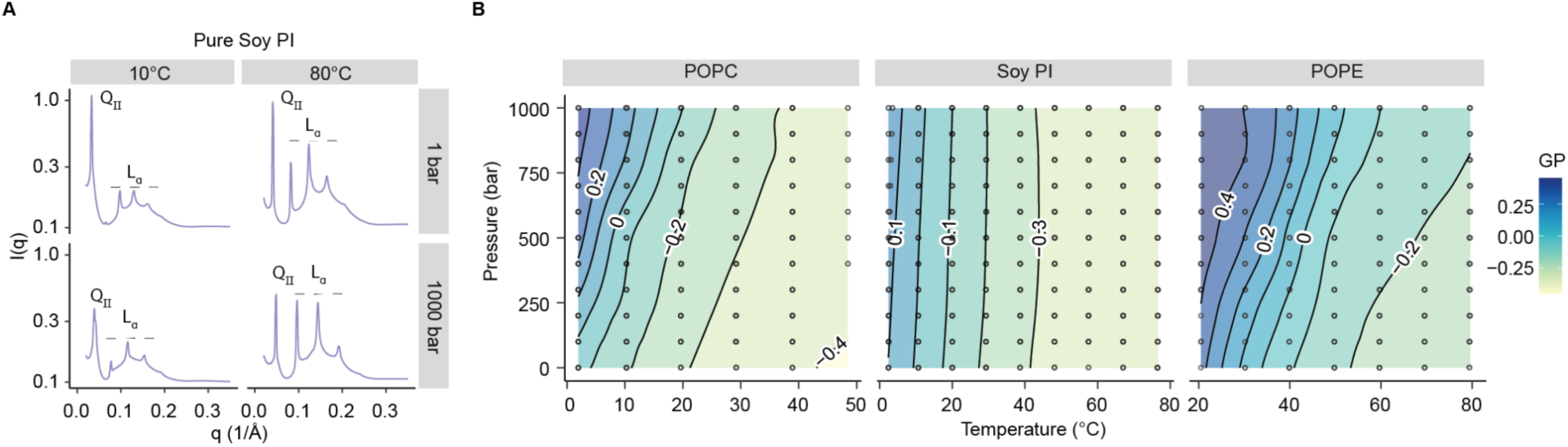
Biophysical properties of Soy PI. (*A*) Representative SAXS profiles taken from suspensions of Soy PI in water at 1 and 80 °C and 1 and 1000 bar, with labels indicating peaks representing the Q_II_ and L_α_ phases. Baselines are offset for clarity. ***(B)*** Pressure-temperature phase diagrams of C-Laurdan GP values of pure POPC, Soy PI, or POPE. GP was determined for temperatures and pressures between 0-80°C and 1-1000 bar. Lines show fitted contours of constant GP values across temperature and pressure, while the scale key represents the coding of GP value for color. Across temperature and pressure,

**Supplementary Figure 4:**
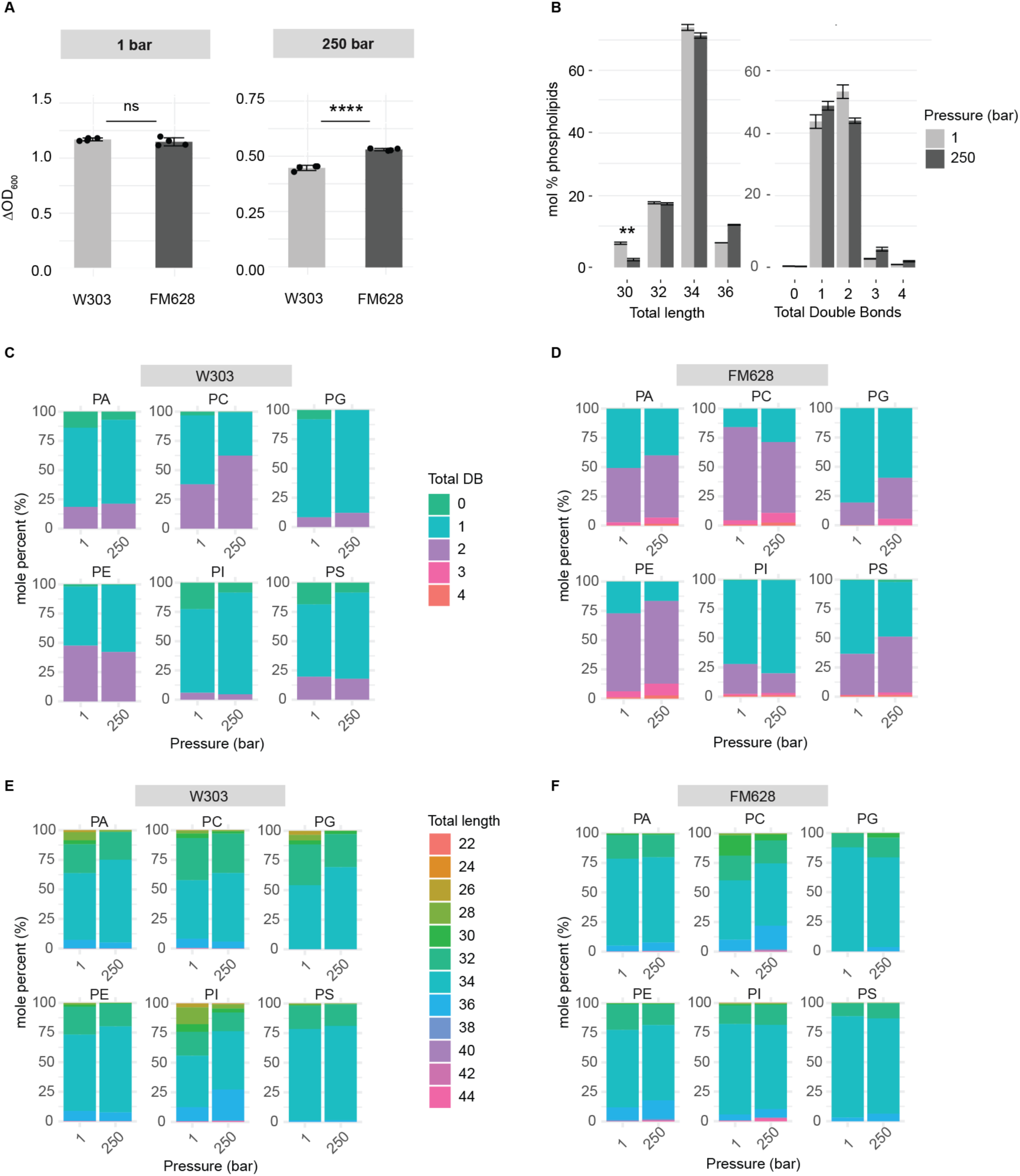
Additional lipidic changes in pressure-grown cells. (***A***) Growth of W303 and FM628 at 1 and 250 bar pressure. Error bars indicate SEM from *N* = 4 independent cultures. **** *P* < 0.0001, unpaired two-tailed test. (***B***) Change in phospholipid chains of FM628 in total length (left) and total double bonds (right). Error bars indicate SEM from *N* = 3 independent cultures. ** *P* < 0.01, unpaired two-tailed test. (***C***) and (***D***): Distribution of total double bonds (DB) by phospholipid class from W303 (C) and FM628 (D) grown at 1 and 250 bar. (***E***) and (***F***): Distribution of total chain length by phospholipid class from W303 (C) and FM628 (D) grown at 1 and 250 bar.

**Supplementary Figure 5:**
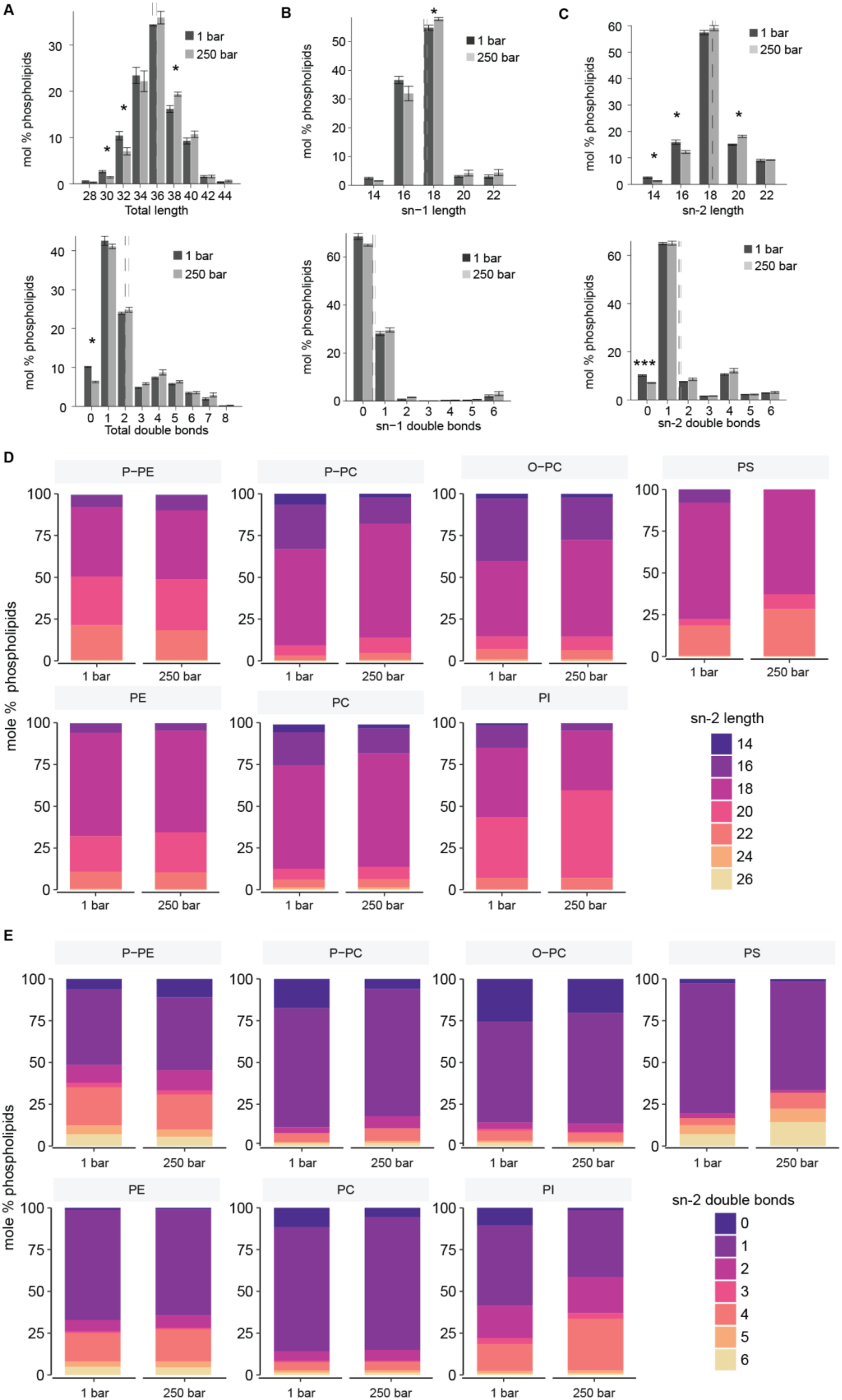
Changes in chain composition of K562 cells grown at 1 and 250 bar pressure. (***A***) Mean total lengths of phospholipid chains in K562 cells incubated at 1 and 250 bar pressure from individual culture triplicates. The mean length of all phospholipids for each pressure is shown with a dashed line. * *P* < 0.05, unpaired two-tailed t-test. (***B***) Mean sn-1 double bonds and length of phospholipid chains in K562 cells incubated at 1 and 250 bar pressure from individual culture triplicates. * *P* < 0.05, unpaired two-tailed t-test. (***C***) Mean sn-2 double bonds and length of phospholipid acyl chains in K562 cells incubated at 1 and 250 bar pressure from individual culture triplicates. * *P* < 0.05, *** *P* < 0.001, unpaired two-tailed t-test. (***D***) Mean sn-2 chain length within each phospholipid class of yeast growth at 1 and 250 bar pressure from individual culture replicates. (***E***) Mean sn-2 chain double bonds within each phospholipid class of yeast growth at 1 and 250 bar pressure from individual culture replicates.

**Supplementary Figure 6:**
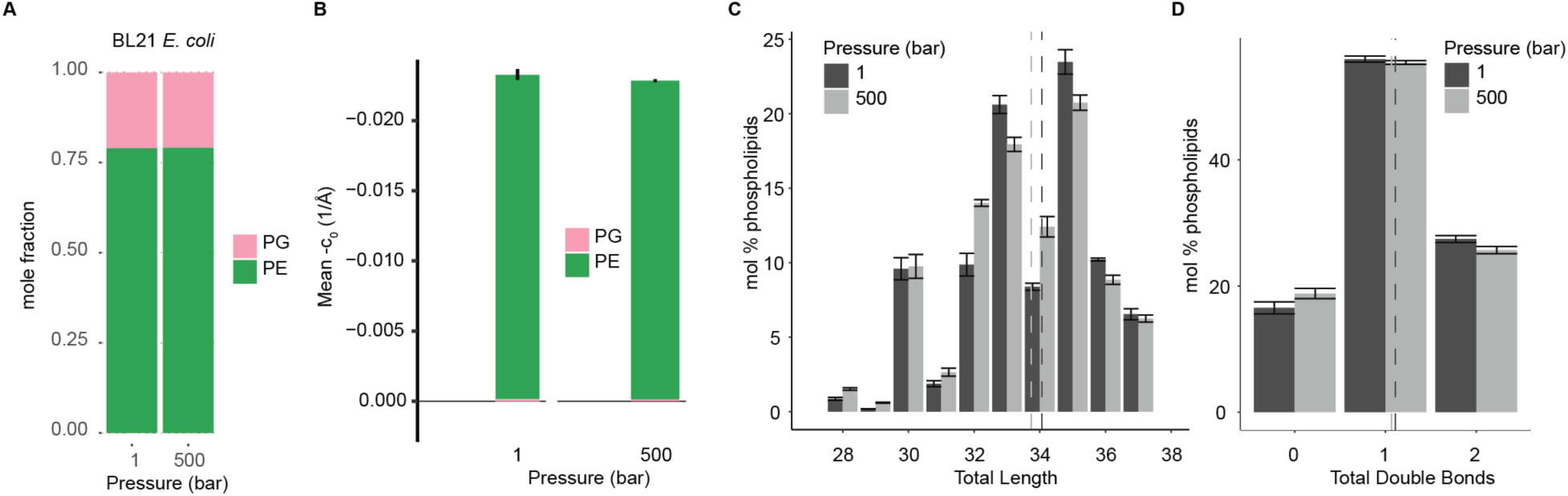
BL21 *E. coli* lipidomics and mean curvature changes from growth at 1 and 500 bar. (***A***) Class composition of phospholipids in *E. coli* grown at 1 and 500 bar pressure. Data is mean mole fraction from individual culture triplicates. (***B***) Mean curvature of phospholipids in *E. coli* grown at 1 and 500 bar pressure from individual culture triplicates. (***C***) Mean total lengths of phospholipid acyl chain composition in *E. coli* grown at 1 and 500 bar pressure from individual culture triplicates. The mean length of all phospholipids for each pressure is shown with a dashed line and mildly decreases at 500 bar. (***D***) Mean total double bonds of phospholipid chain composition in *E. coli* grown at 1 and 500 bar pressure from individual culture triplicates. The mean double bonds of all phospholipids for each pressure is shown with a dashed line and is unchanged with pressure.

**Table 1:** Yeast strains used in this study.

| Name | Genotype | Parental strain | Reference |
| --- | --- | --- | --- |
| FM628 | *Lachancea kluyveri* | FM628 | Maitreya Dunham |
| W303a | *MATa ura3-52 trp1Δ2*<br>*leu2-3_112 his3-11*<br>*ade2-1 can1-100* | W303a | ATCC |
| *pem2Δ* | W303a, *pem2Δ::HIS3* | W303a | This study |
| *psd1Δ* | W303a, *psd1Δ::HIS3* | W303a | This study |

